# Unoccupied aerial system enabled functional modeling of maize (*Zea mays* L.) height reveals dynamic expression of loci associated to temporal growth

**DOI:** 10.1101/848531

**Authors:** Steven L. Anderson, Seth C. Murray, Yuanyuan Chen, Lonesome Malambo, Anjin Chang, Sorin Popescu, Dale Cope, Jinha Jung

**Affiliations:** Department of Soil and Crop Sciences, Texas A&M University, College Station, TX 77843; Department of Ecosystem Science and Management, Texas A&M University, College Station, TX, 77843; School of Engineering and Computer Sciences, Texas A&M University - Corpus Christi, Corpus Christi, TX 78412; Department of Mechanical Engineering, Texas A&M University, College Station, TX 77843

## Abstract

Unoccupied aerial systems (UAS) were used to phenotype growth trajectories of inbred maize populations under field conditions. Three recombinant inbred line populations were surveyed on a weekly basis collecting RGB images across two irrigation regimens (irrigated and non-irrigated/rain fed). Plant height, estimated by the 95^th^ percentile (P95) height from UAS generated 3D point clouds, exceeded 70% correlation to manual ground truth measurements and 51% of experimental variance was explained by genetics. The Weibull sigmoidal function accurately modeled plant growth (R^2^: >99%; RMSE: < 4 cm) from P95 genetic means. The mean asymptote was strongly correlated (r^2^=0.66-0.77) with terminal plant height. Maximum absolute growth rates (mm d^-1^) were weakly correlated to height and flowering time. The average inflection point ranged from 57 to 60 days after sowing (DAS) and was correlated with flowering time (r^2^=0.45-0.68). Functional growth parameters (asymptote, inflection point, growth rate) alone identified 34 genetic loci, each explaining 3 to 15% of total genetic variation. Plant height was estimated at one-day intervals to 85 DAS, identifying 58 unique temporal quantitative trait loci (QTL) locations. Genomic hotspots on chromosome 1 and 3 indicated chromosomal regions associated with functional growth trajectories influencing flowering time, growth rate, and terminal growth. Temporal QTL demonstrated unique dynamic expression patterns not observable previously, no QTL were significantly expressed throughout the entire growing season. UAS technologies improved phenotypic selection accuracy and permitted monitoring traits on a temporal scale previously infeasible using manual measurements, furthering understanding of crop development and biological trajectories.

**Author summary:** Unoccupied aerial systems (UAS) now can provide high throughput phenotyping to functionally model plant growth and explore genetic loci underlying temporal expression of dynamic phenotypes, specifically plant height. Efficient integration of temporal phenotyping via UAS, will improve the scientific understanding of dynamic, quantitative traits and developmental trajectories of important agronomic crops, leading to new understanding of plant biology. Here we present, for the first time, the dynamic nature of quantitative trait loci (QTL) over time under field conditions. To our knowledge, this is first empirical study to expand beyond selective developmental time points, evaluating functional and temporal QTL expression in maize (*Zea mays* L.) throughout a growing season within a field-based environment.

## Introduction

Phenotypic characterization of agricultural plant populations has lagged in scale, density, and accuracy when compared with genomic data [1]. Due to resource demands of labor and time-sensitive components in conventional phenotyping, most manually measured traits are acquired at only one time point in the growing season and constrained in the number of samples. This creates a limited scope of biological understanding when associating genomic information with the underlying traits of interest through plant development [2]. Advances in technologies including computer vision, robotics, remote sensing, and unoccupied vehicles have facilitated the development of high-throughput phenotyping (HTP) platforms which can minimize phenotypic bottlenecks [3, 4].

Implementation of HTP systems provides the ability to collect temporal phenotypic measurements on large representative populations within field settings, to understand how individuals interact with their environments [3, 5, 6]. Unoccupied aerial systems (UAS) are especially useful to increase the size of populations and field studies investigated, collecting RGB images, and reconstructing three dimensional representations of field crop trials using structure from motion methodology [6–15]. UAS height estimates of maize have previously been validated using correlations to traditional manual measurements and evidence of equivalent or greater phenotypic variation partitioned to genetic factors [7, 9, 15, 16]. To our knowledge the majority of reported field based phenotyping of maize with HTP platforms has focused on hybrid trials [6-9, 15, 17-19] but, limited reports have been published on the evaluation of inbred trials [20–22], specifically genetic mapping populations. Inbred lines in maize are substantially shorter and have less biomass than hybrids, lacking heterosis.

Maize height is important as a physiological and a highly heritable agronomic trait [23–26] commonly collected due to its ease of measurement, agronomic importance, and correlation to hybrid grain yield in some environment and management scenarios [15, 27–32]. Manually measured plant height is commonly collected after reproductive maturity as the distance from the ground to the tip of the tassel, flag leaf, or peduncle. The genetic architecture of plant height in maize has been determined to fit an infinitesimal model (i.e. very large numbers of small additive effect loci) with some large effect loci likely fixed during domestication and early selection [23]. Functional genetic variation in terminal plant height has been shown to be controlled through hormones; mutations within the (i) gibberellin biosynthesis pathways [33] and crosstalk with other phytohormomes including: (ii) auxin [34] and (iii) brassinosteriods [35–38]. Hormones are well known to fluctuate throughout plant growth, responding to environmental and developmental stimuli [39–42]. Traditional QTL studies using phenotypic data at a single terminal (end of season) time point can only represent accumulated effects, ignoring the dynamic nature of many agronomically important traits which, like hormones, change and can be identified as functions of time [43]. To our knowledge, Wang, Zhang (20) is the only reported field based temporal association study in maize using UAS.

Plant height is an ideal phenotype to explore the temporal patterns of QTL expression in maize. Using UAS, we evaluated three recombinant inbred line (RIL) linkage mapping populations under field conditions and captured the dynamic growth patterns of plant height across these maize inbreds. The objectives of this study were to: (i) evaluate UAS procedures developed for hybrids to estimate heights within inbred maize populations; (ii) model and compare growth patterns across genetic populations; (iii) evaluate temporal patterns of QTL expression through the growing season, and (iv) evaluate the temporal expression patterns for previously reported QTL.

## Results and Discussion

### UAS surveys and image processing quality

A total of 18 and 11 flights were conducted over the bi-parental mapping populations using the DJI Phantom 3 Pro and Tuffwing UAV Mapper, respectively (S1 Table). Early season DJI Phantom 3 Pro data collection prior to 35 DAS resulted in limited to no plant structure reconstructed within the 3D point clouds, indicating that higher resolution imaging would be necessary to reconstruct early season plant structure. Out of 29 flights, 16 were observed to be of high quality while only eight flight dates (35, 43, 57, 62, 65, 69, 100, and 117 DAS) conformed to statistical quality tests (S1 File) and were used for the remainder of this study (Fig 1; S2 Table).

**Fig 1.**
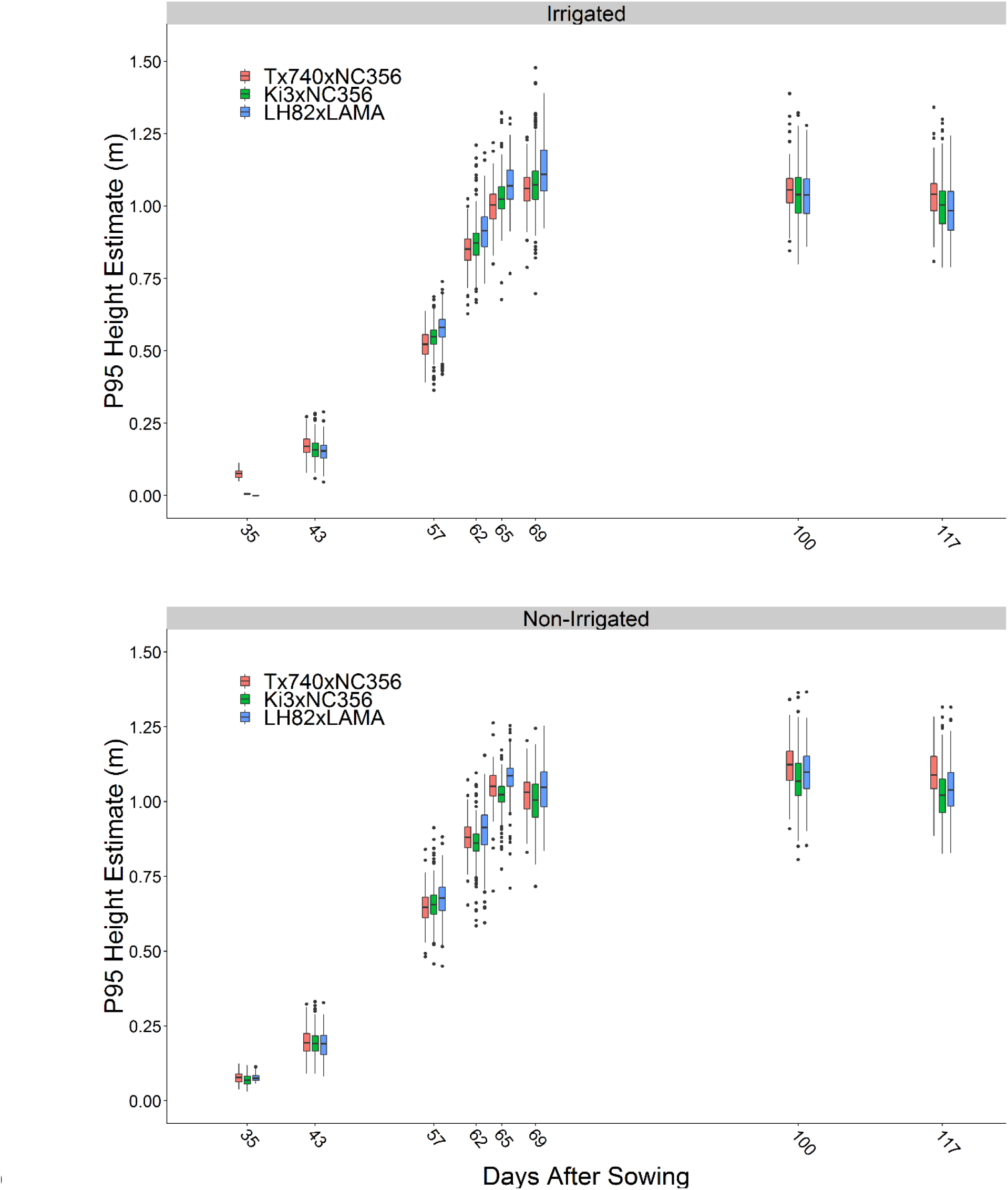
UAS P95 height estimates summarized by flight date. Although the three populations were genetically diverse, the mean growth patterns behaved similarly. Little differentiation could be seen early in the season between genotypes, where the measurement error may have been smaller that genotypic differences, as the plants reached their peak height and flowered, height differences became much greater.

### Genetic variance decomposition

Variance component decomposition demonstrated total phenotypic variance increased throughout the growing season for all inbred populations (Fig 2, black circles), as has been found in hybrid trials [7, 15]. UAS phenotypic variance for height did not exceed manual, terminal plant height (PHT_TRML_, Fig 2 M bar). Genetic variance averaged 51% (excluding 35 DAS) over the season fluctuating from flight to flight, but generally increasing until reaching a terminal height plateau. The proportion of variance attributed to genetics of plant height (PHT_TRML_), as measured from the ground to the tip of the tassel, were numerically greater (irrigated: 62 ± 3%; non-irrigated: 52 ± 3), but not statistically (α=0.05) different from across UAS genetic variance.

**Fig 2.**
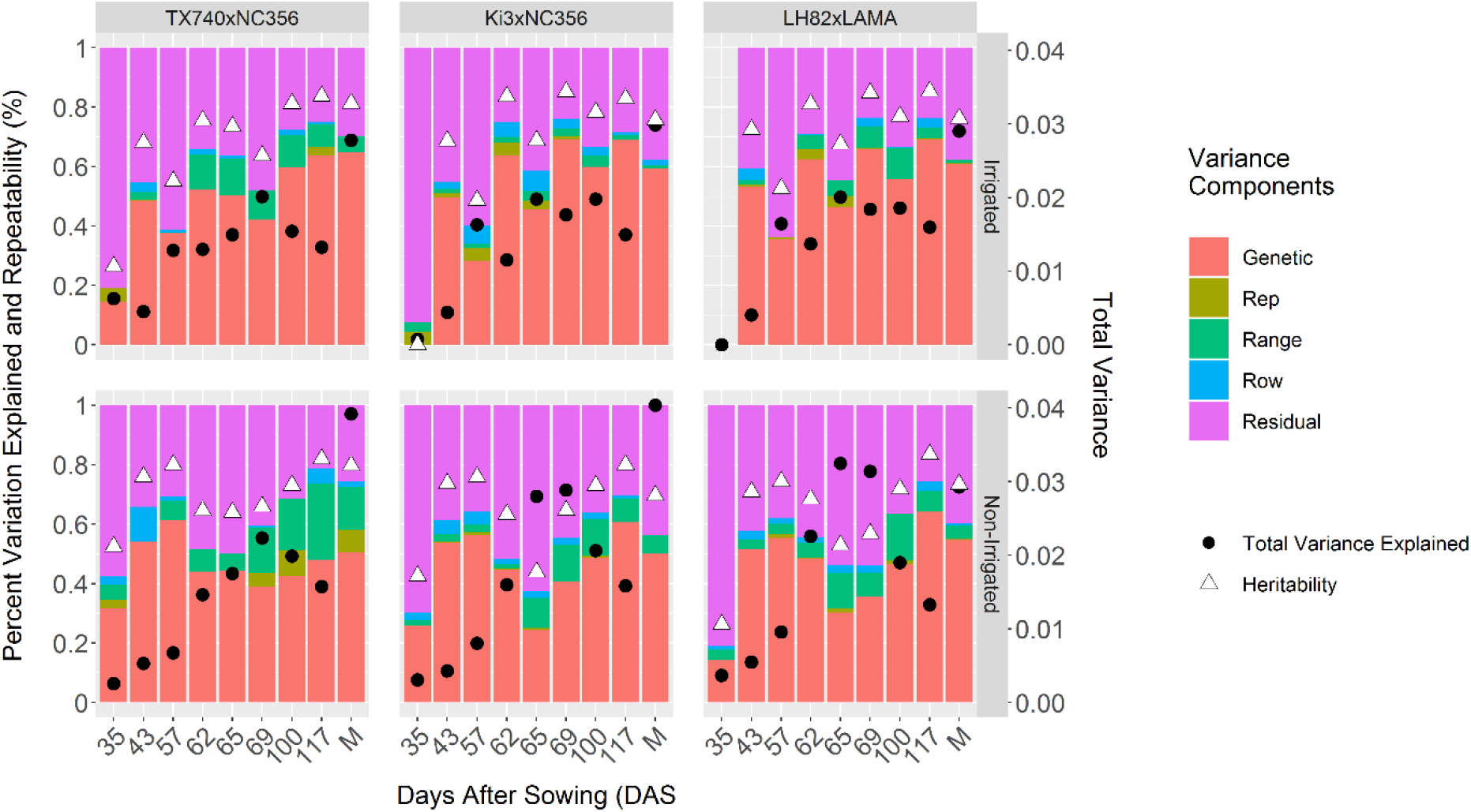
Variance component decomposition of UAS P95 height estimates. The percent variation explained in the model of Eq. 1 for individual UAS surveys of three RIL populations showed that genetic and residual (error) variation, were the main drivers of variability observed. Total variance (black circles) increased as the plants grew over later flight dates and was higher for manual (M) than UAS measurements. That the percent variance measures and heritability were similar for M and UAS suggests that UAS compressed all variance sources similarly

### Sigmoidal modeling of UAS height estimates

The Weibull function, a sigmoidal growth function, modeled maize inbred temporal growth (mean R^2^>0.99, RMSE ranging from 2.4 to 3.7 cm) across all populations and environment (Fig 3). Significant differences in asymptote, the maximum height, were only found between Tx740xNC356 (1.10 m) and LH82xLAMA (1.08 m) with a 2 cm difference in means under irrigation. LH82 [44] is the earliest to flower and shortest of the inbred lines adaptable to these environments and had among the lowest asymptote and inflection point, but moderate growth rate.

**Fig 3.**
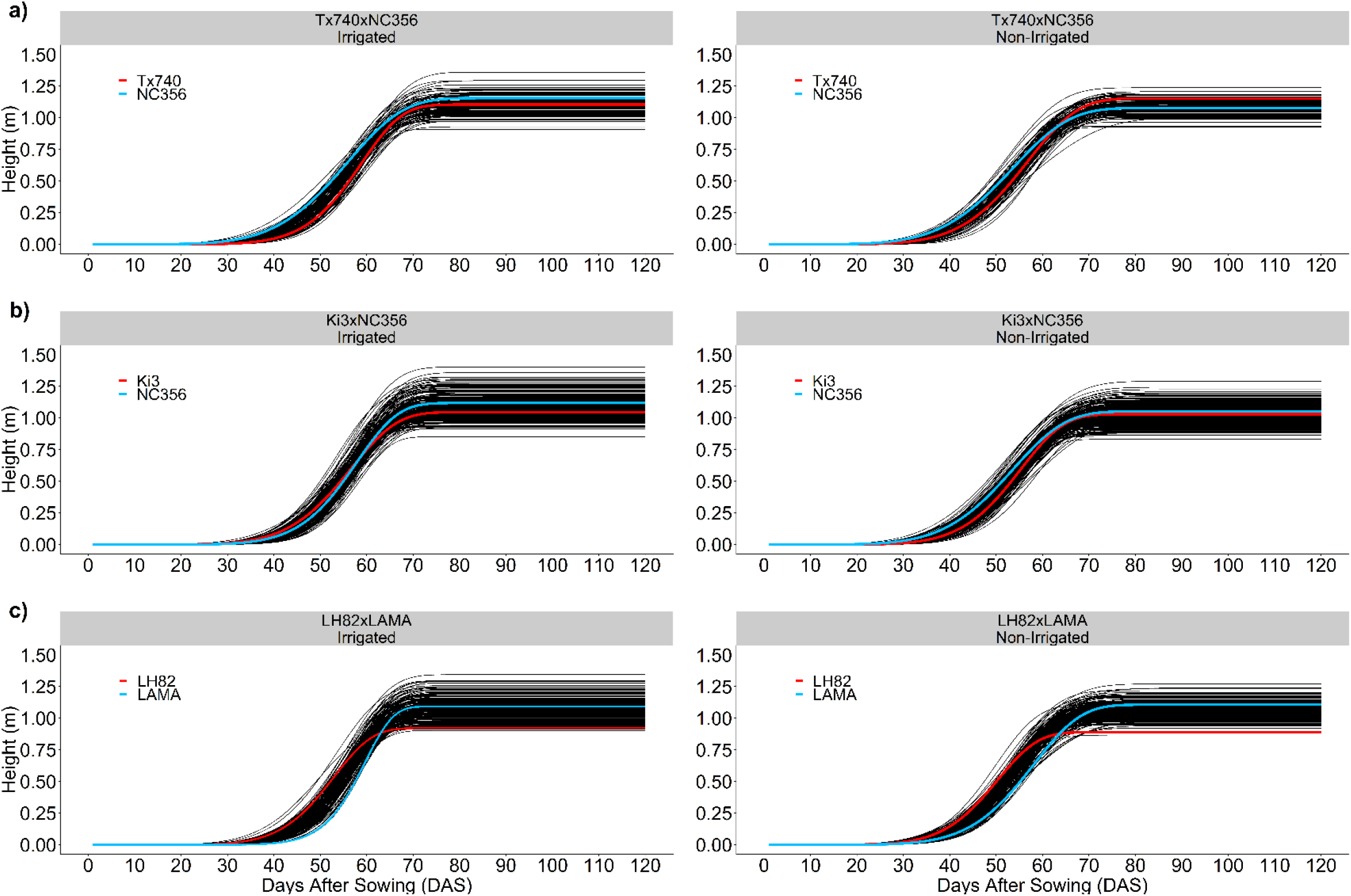
Nonlinear Weibull functional modeling of growth trajectories. Sigmoidal curves based off the Weibull function (Eq. [3]) effectively modeled the growth of each entry. For each population the female parent (red line) and the male parent (blue line) crossed over demonstrating that early season height was not predictable by standard manual terminal height measurements.

In comparison, PHT_TMRL_ was significantly different across populations (1.66, 1.59, and 1.57 m) under irrigated conditions for Tx740xNC356, Ki3xNC356, and LH82xLAMA, respectively. The reduced means of the asymptote demonstrate the inherent biases of UAS estimation of plant height compared with manual measurements [8, 12, 13, 15]; ∼0.5 m underestimate of height has been documented in past studies of hybrid maize at flight altitudes of 120 m [15]. The average difference in height estimates increased by ∼5 and ∼10 cm when compared to P99 and P100 point cloud estimates, indicating that the reduction was not caused solely by the lower percentile, P95. The combination of flight altitude and reduced plant canopy density of the inbreds likely biased the UAS towards shorter estimates. Biases aside, numerical rankings between asymptote and PHT_TRML_ were correctly consistent in ranking Tx740xNC356, Ki3xNC356, and LH82xLAMA population means from tallest to shortest and Pearson correlations (r) (Irrigated: 0.77, 0.74, and 0.74; Non-Irrigated: 0.66, 0.72, and 0.74; S1-S3 Fig) indicated highly significant (α=0.05), positive linear correlations between UAS asymptotes estimates and PHT_TRML_ measurements.

The inflection point of the Weibull model is biologically important to identify the DAS in which maximum AGR is occurring; this point has been shown to be highly correlated to flowering time in hybrid trials [15]. Significant differences were found between each population’s mean for inflection point (58.6, 58.0, and 57.5 d for Tx740xNC356, Ki3xNC356, and LH82xLAMA) within the irrigated trial (Fig 4c). Abiotic stress related to water limitations in non-irrigated trials delayed the inflection point by two days on average across the populations. Inflection point had low positive correlations to PHT_TRML_ (Irrigated: 0.30, 0.27, and 0.34; Non-Irrigated: 0.02, 0.22, and 0.24; S1-S3 Fig) but high correlations to flowering time (DTA/DTS) (Irrigated: 0.60/0.45, 0.59/0.58, and 0.64/0.59; Non-Irrigated: 0.61/0.56, 0.55/0.53, and 0.68/0.66; S1-S3 Fig). PHT P95 estimates were negatively correlated (r= −0.74:-0.50) to inflection points during the early season but gradually progressed toward a positive correlation ∼10 days after the mean inflection point (S1-S3 Fig). Later inflection points had extended vegetative growth periods leading to taller plants, indicating the possibility of pleiotropic QTL for both flowering time and growth rate across the functional curve parameters. Because correlation was high but imperfect, tall genotypes with earlier inflection points could indicate better fitness in stressful environments, as these plants reach their terminal height quickly without regard to environmental stresses.

**Fig 4.**
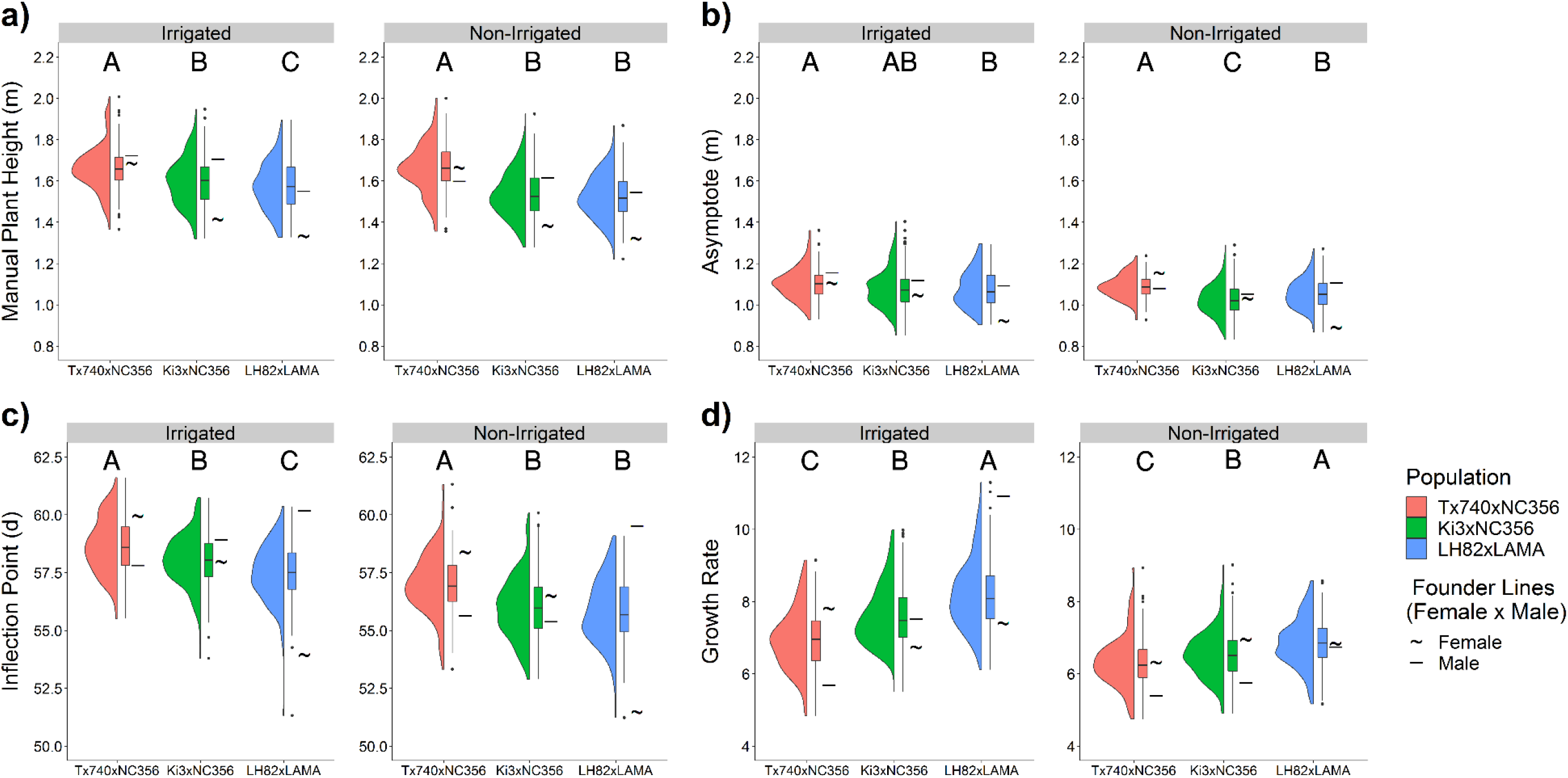
Distribution of Weibull functional parameters. Entry BLUPS of **[a]** manual terminal plant height, **[b]** Weibull asymptote, **[c]** Weibull inflection point, and **[d]** Weibull growth rate for each mapping population demonstrated variability both within and between these populations with substantial transgressive segregation in most cases. Letters above define significant differences in means at α=0.05.

The growth rate parameter, influencing the steepness of the Weibull curve, significantly differed (α=0.05) in its means across the populations in both environments (Irrigated: 6.9, 7.6, and 8.2; Non-irrigated: 6.3, 6.5, and 6.8). The first derivative of the Weibull function (Eq. 4), the absolute growth rate (AGR), calculated at the inflection point (x=x_0_) equals the maximum AGR. The maximum AGR occurred ∼50-60 DAS, which was shortly before flowering, in this period cells are both dividing and elongating within the internodes above the ear node [45–47]. Significant differences were found in the maximum AGR across populations within the irrigated trial (48, 52, and 56 mm d^-1^) and LH82xLAMA was 3 mm d^-1^ greater than the other populations in the non-irrigated trial despite being the shortest population overall. A reduction in AGR was observed within the non-irrigated trial (4, 7, and 8 mm d^-1^ Tx740xNC356, Ki3xNC356, and LH82xLAMA, respectively) likely due to water stress during this period [48]. Overall this demonstrated that the AGR had heritable genetic diversity and was phenotypically plastic in response to different environmental conditions.

### QTL mapping

#### Manual terminal height associations

Nine QTL were identified for PHTTRML across the three populations and two environments (S3 Table) each explaining 5.1 to 9.4% of genetic variance. All PHTTRML associations had additive effects of ∼3 cm (S3 Table). One region was identified across two populations q1_172 (LH82xLAMA; irrigated) and q1_176 (Tx740xNC356; non-irrigated), localizing to the 280 to 284 Mbs region of chromosome 1. We identified a single genomic region, 98 to 128 Mbs on chromosome 2 that co-localized within the same genetic background (Ki3xNC356) across different environmental treatments (q2_70 irrigated and q2_69 non-irrigated). The limited co-localization of QTLs across bi-parental populations is part of the difficulty of identifying genomic regions that can be utilized in genetic backgrounds beyond those in which they were discovered [49, 50]. This also demonstrated the lack of statistical power in the smaller of the three populations Tx740xNC356 (n=110). It has been empirically shown that population size is the most critical factor in QTL linkage mapping [24].

#### Functional parameter associations

UAS estimates in modeling temporal growth of maize can identify dynamic QTL [51]. Analysis of QTLs using the three functional parameters of the Weibull curve as phenotypes identified 13, 9, and 12 significant marker associations with the asymptote, growth rate, and inflection point, respectively (S5 Table). Asymptote QTLs explained genetic variation ranging from 3.4% to 14.3% with additive effects ranging from 2 to 5 cm, consistent with PHTTRML. High correlations between asymptote and PHTTRML indicated that similar QTL would likely be detected using both traits. Two PHTTRML QTLs, q1_172 LH82xLAMA (irrigated) and q1_176 Tx740xNC356 (non-irrigated), co-localized with an asymptote QTL, q1_173 of LH82xLAMA (irrigated) (Fig 5; S3 Table). Additional co-localizations were found between q6_67 Tx740xNC356 (irrigated) asymptote and q6_62 Ki3xNC356 (irrigated) PHTTRML, as well as, q8_10 LH82xLAMA (non-irrigated) asymptote with q8_14 Ki3xNC356 (irrigated) PHTTRML and q8_12 Ki3xNC356 (non-irrigated) PHTTRML.

**Fig 5.**
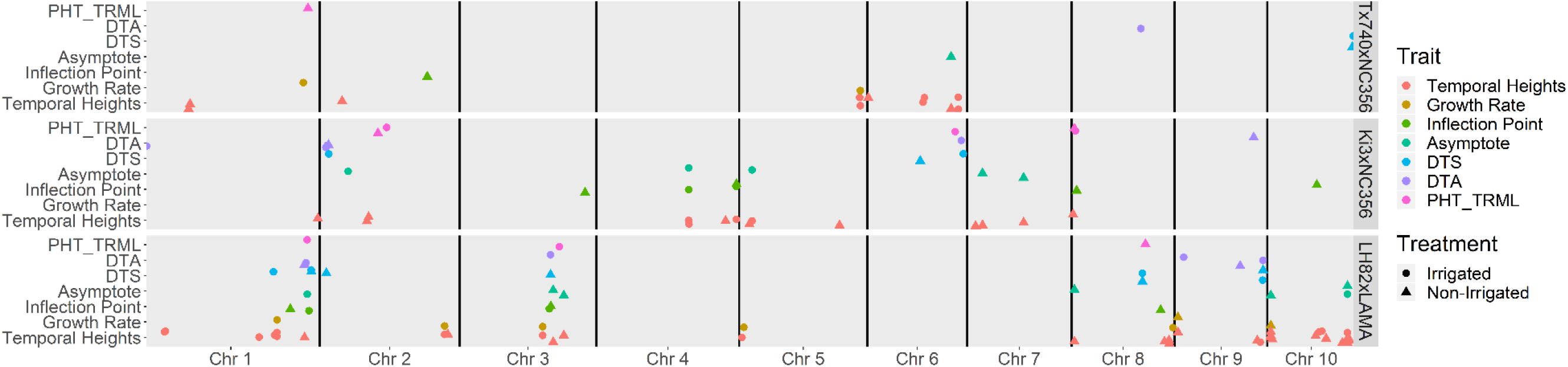
Col-localization of agronomic and functional growth QTL associations. Significant QTL co-localized across agronomic traits (PHT_TRML: Manual, terminal plant height; DTA: Days to anthesis; DTS: Days to silking), functional growth parameters (asymptote, inflection point, growth rate) and temporal height estimates from the Weibull curves. Temporal expression of all height QTL can be visualized in **S4 Fig**.

The seven growth rate QTL each explained 5.6 to 15% of the genotypic variance with additive effects ranging from 0.2 to 0.3 DAS^-1^ (S5 Table). Inflection point QTL each explained 4.3 to 13% of the genotypic variance with additive effects ranging from 0.2 to 0.5 d (S5 Table). Irrigated Ki3xNC356 trial *q*4_61 and irrigated LH82xLAMA *q*1_173/*q*1_176 were associated with inflection point and asymptote, while non-irrigated LH82xLAMA *q*10_20 was associated with inflection point and growth rate (S5 Table). The co-localization of QTL associated with multiple parameters of the sigmoidal growth function indicated these regions more than others may have an effect on defining the overall developmental trajectory of maize height. The limited number of co-localizations demonstrates these traits are both genetically variable and highly plastic with the environment.

Multiple QTL were identified within the LH82xLAMA trials for PHT_TRML_, asymptote, inflection point, and flowering time (DTA/DTS; S4 Table) within the 273 to 287 Mbs region of chromosome 1 and the 140 to 176 Mbs region of chromosome 3 (Fig 5). The QTL region of chromosome 3 harbors ZmMADS69 (GRMZM2G171650; Chr3: 158979321..159007265), a regulator of flowering time with pleiotropic effects on plant height. ZmMADS69 has higher expression levels in temperate compared to tropical germplasm, leading to significant detection in temperate by tropical crosses [52], such as LH82xLAMA among others [23, 24, 53]. The identified region on chromosome 1 contained the viviparous8 (vp8; GRMZM2G010353; Chr1: 286390345..286398537) locus which exhibits dwarfism due to reduced cell proliferation [54]. Both loci may be deterministic QTL (dQTL) because the differential allelic variation affected the whole growth process [55] and was unaffected by environmental stimuli; ZmMADS69 effect was not influenced by day length [52] and vp8 exhibited normal plant hormone response [54]. These results coupled with basic biological understanding indicated that allelic changes in loci can have a fundamental impacts on the functional growth trajectory of maize, in contrast to the small shift in phenotypic expression of a single trait It is therefore understandable that these two “major genes” have been previously identified and described in multiple studies, while the smaller and ephemeral effect loci are mostly unknown.

#### Temporal QTL expression

In addition to detecting QTL for the three parameters of the Weibull function, 58 QTL were also detected using individuals’ daily heights from 20 to 85 DAS predicted using the Weibull function. Between 4 and 20 unique QTLs were identified, based on peak position (Fig 5; S6 Table). Comparison of mean physical distance of the flanking markers for each the 58 unique QTLs demonstrated 23 QTLs were within 1 Mbp of a plant height candidate gene and an additional 18 QTL were less than 5 Mbp from a candidate gene. Most of the 58 unique QTLs demonstrated a very dynamic nature of QTL affecting plant height throughout the growing season. For example, q5_119 in the irrigated Tx740xNC356 trial, was detected from 22 to 62 DAS explaining 21% of the genetic variation at 54 DAS (Fig 6a; S6 Table). In comparison, q5_35 of irrigated Tx740xNC356 trial was detected from 66 to 74 DAS explaining 11% of the genetic variation at 67 DAS (Fig 6a). Temporal QTL expression was different for each population across environmental treatments (i.e. irrigation) demonstrating differential genomic localization while maintaining similarities in temporal expression. Specifically, within the Tx740xNC356 population, both irrigation regimens (i.e. environments) have a temporally broad QTL (q5_119 irrigated and q2_55 non-irrigated) prior to inflection point (∼58 DAS), followed by QTLs detected at shorter temporal intervals after the inflection point and may relate to the elongation of specific internode groupings [45–47]. Additionally, trends in QTL temporal expression between populations exhibited unique temporal expression patterns. For example, Tx740xNC356 exhibited QTLs prior to the inflection point at early growth stages, whereas Ki3xNC356 exhibited no detectable QTLs until ∼50 DAS. Low phenotypic variation could be the cause, as could, greater numbers of smaller effect loci, towards an infinitesimal model, that would also be hard to detect.

**Fig 6.**
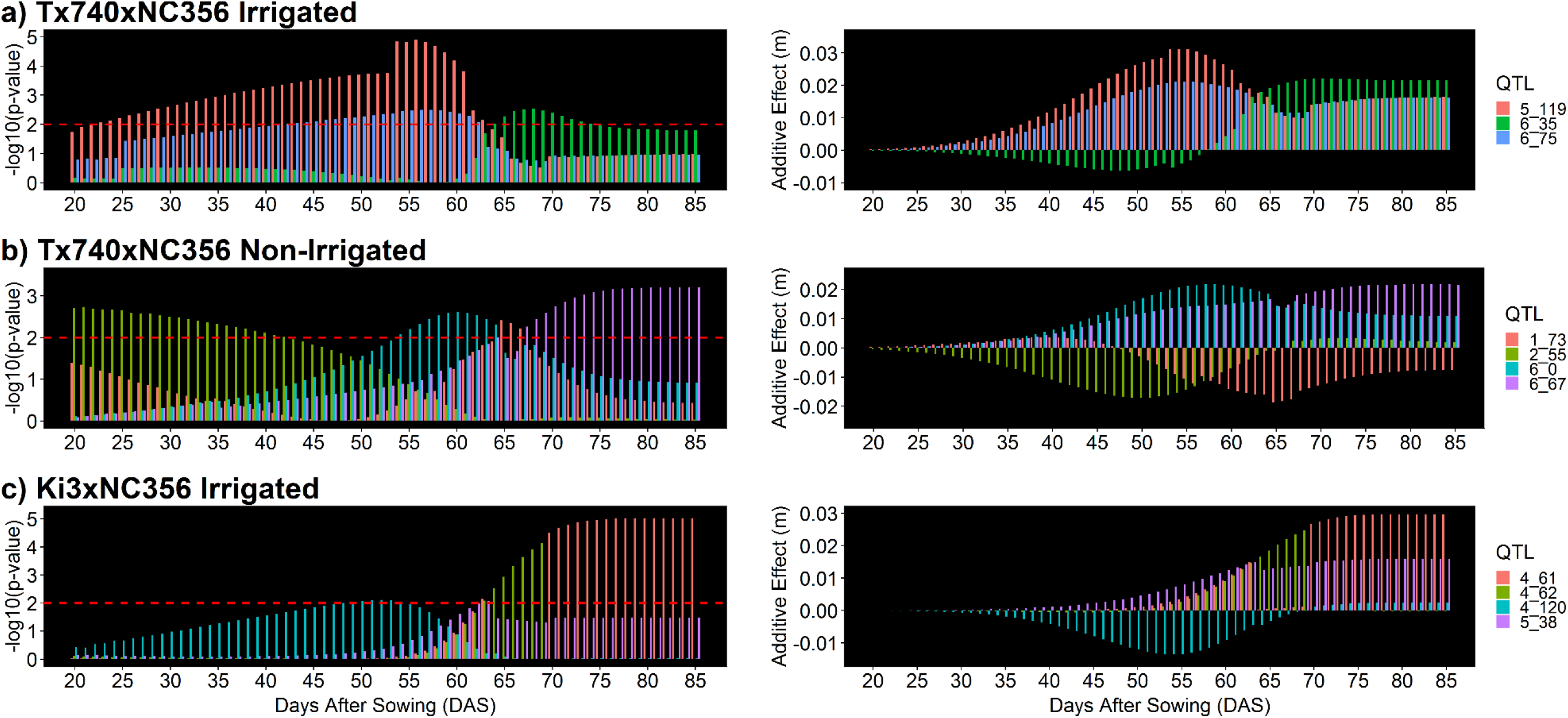
Visualization of temporal trends in QTL expression. Temporal trends in QTL expression were observed in all QTL across populations and environments. Most QTL were under the significance threshold (left side) of LOD=2 (red dashed line) at some point during the growing season; however the smaller additive effects (right side) during these periods would not have been expected to be declared a QTL.

Identified QTLs demonstrated dynamic trends in additive phenotypic effects (Fig 6, right side). In general, these results show that the additive effects found at the peak significance DAS of a temporal QTL is a result of the cumulative effect of a gradual increase in the effect size of each genomic region (Fig 6). QTLs with peak expression early within the season had significantly smaller additive effect estimates than at later points in the growing season; due to reduced overall variation across individuals in the population at early growth stages (e.g. Fig 6b *q*2_55). Some QTL effects (Fig 6b *q*6_0) also appeared to lose their association throughout season, however it is likely due to their effects statistically being diluted by new QTL that become significant (Fig 6b *q*6_67). While most individual QTL maintained their directional effect (Fig 6a; q5_119 and *q*6_75), some surprisingly switched effect directions within the growing season (q6_35). Understanding the biological basis of this switching phenomena would be both interesting and important for optimizing plant growth. Caution should be used to interpret all of these QTL as loci that functionally affect height and plant growth rather than height QTL per se; loci affecting rooting, plant health or photoperiod sensitivity all could impact plant measured height.

Analysis across the entire linkage map demonstrated that directional changes in additive effect size were present during the growing season (S4 and S5 Fig). Within marker assisted selection protocols, targeting consistent directional effects may result in greater gains than those of temporal bi-directional effects. Before additional work is conducted temporal effect size should first be validated through near isogenic lines across genetic backgrounds or in heterogeneous inbred families [56]. However, we speculate that the temporal trend of the effect size, like many QTL effects remains dependent on the genetic background, abiotic, and biotic interactions experienced in each environment, and this G x E interaction. If temporal shifts in directional effect are valid and not due to over inflations via false positives and limited population size; statistical models accounting for directional effect shifts will be necessary to incorporate temporal datasets of dynamic, quantitative traits within prediction modeling approaches to plant breeding, such as genomic selection.

## Materials and methods

### Germplasm material and experimental design

Three bi-parental mapping populations were developed from breeding lines segregating for loci discovered in an earlier genome wide association study [28, 57] of hybrids for height and grain yield. The recombinant inbred line (RIL) progeny were derived from the crosses of Tx740xNC356 (tropical/tropical; 110 RILs), Ki3xNC356 (tropical/tropical; 239 RILs) and LH82xLAMA-YC (temperate/tropical; 178 RILs). Tx740 (LAMA2002-12-1-B-B-B) [58] is a parent in the “LAMA” inbred line (pedigree [((LAMA2002-12-1-B-B-B-B/LAMA2002-1-5-B-B-B-B)-3-2-B-1-B3-B]) and these two lines would be expected to share 50% of their genome. In 2018, the mapping populations were planted in a randomized complete block design (RCBD) with two replications across two environments (irrigated and non-irrigated) having dimensions of 0.76 m row spacing, and 3.81 m plot lengths.

### Unoccupied aerial system image collection

Two platforms were used, a rotary wing and a fixed wing UAV, to collect RGB data. For the rotary wing, a DJI Phantom 3 Professional with a 12-megapixel DJI FC300X camera was flown at an altitude of 25 m with to 80% forward and side image overlap. Fixed wing images were collected using a Tuffwing UAV Mapper (http://www.tuffwing.com) equipped with a 24-megapixel Sony a6000 RGB camera. Fixed wing surveys were conducted at a 120 m altitude with 80% image overlap. A total of 19 DJI Phantom 3 Professional flights were conducted throughout the growing season, while 11 Tuffwing UAV Mapper flights (starting 05/17/2018) were conducted after early season to mechanical setbacks of the Tuffwing (S2 Table). After QC/QA, a total of 16 flights were used for height estimates based on quality of the processed orthomosaic images.

All of the Tuffwing flights were processed in Agisoft PhotoScan [59], while the majority of the DJI Phantom flights were processed in Pix4Dmapper [60], based on collaborators comfort and preference with the associated software. In general, these software packages are equivalent and used to identify common features (tie points) across images followed by triangulation and distortion adjustment optimization to generate densified 3D point clouds, DSM, and orthomosaic images. Height estimates were extracted from the three dimensional point clouds following the procedures of [15]. In short, the ground points were identified from the point cloud using the hierarchical robust interpolation algorithm within FUSION/LDV. Identified ground points were used to interpolate the digital elevation model, followed by subtracting the DEM from the original point cloud to produce the canopy surface model. The plot level polygon shapefiles were created using the R/UAStools::plotshpcreate [15] function in R and the 95^th^ percentile height estimates were extracted for each experimental plot.

### Statistical Inference

#### Variance component estimates and heritability

From the extracted canopy height metrics (P95), we fit mixed linear models utilizing residual maximum likelihood (REML) in JMP version 14.0.0 [61] to define best linear unbiased predictors (BLUPs) of the inbreds by their entry number. Models were fit on a per flight date basis. The individual mapping populations were evaluated as a randomized complete block design (RCBD, Eq. 1) including spatial regression (range and row [furrow irrigation runs down rows], this is called row and column, respectively, where furrow irrigation is not used).

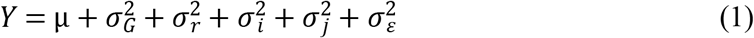

with terms entry 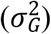, replicate 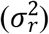 range 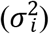, row 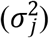and residual error 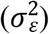.

Broad sense heritability (H^2^) estimates were calculated on an entry means basis (Eq.2).

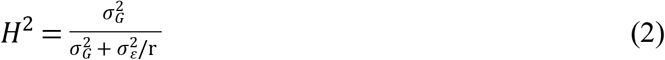

Within each environment, H^2^ estimates were calculated for each population separately while including replicates (r) for each of the UAS flight dates.

#### Nonlinear function

The three parameter Weibull sigmoid growth model (Eq. 3) was used to summarize the

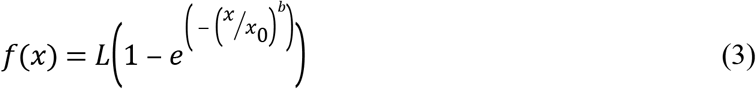

height as a function of DAS (x) with the asymptote (L), inflection point (x_0_), and the growth rate (b) of the fitted curve. The asymptote (L; m) is maximum value of the curve which represents maximum/terminal plant height (PHT_TRML_). The inflection point (x_0_; DAS) indicates the DAS where the slope of the logarithmic phase is at its absolute maximum. The growth rate (b) is an empirical constant which defines the shape of the curve and relates to the absolute growth rate (Eq. 4; m d^-1^) when x=x_0_. Sigmoidal curves were fit using the Fit Curve tool in JMP 14 and

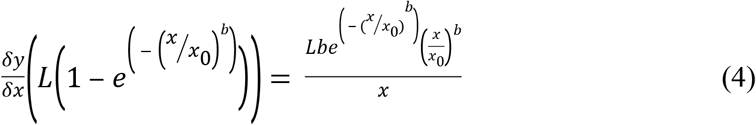

parameters were estimated on an entry basis utilizing the extracted BLUPs or the individual environment REML models described above. Significance of the functional parameters were evaluated using the chi squared (X^2^) test (α = 0.05, df = 1) to identify logistical curves with poor fits to UAS height estimates, these were subsequently removed from future analysis. Using the associated Weibull functional parameters, height estimates were imputed on one day intervals (1 to 85 DAS) for each inbred entry in their associated environments.

### Genotyping and linkage map construction

The genotyping was described in Chen, Murray (62), and is paraphrased here. Genotype samples were collected from F_3:4_ seedlings grown under greenhouse conditions, where eight samples were bulked per genotype. The CTAB method [63] was used to extract DNA and samples were sent to AgReliant Genetics LLC, where they were genotyped by Infinium® assays for 17,444 single nucleotide polymorphisms (SNPs). The linkage groups and physical locations were provided with the SNP chip of which 716 markers locations were unknown or withheld due to intellectual property rights, resulting in 17,019 SNPs with known reference locations (B73 RefGEN_v3).

Individuals with >10% missing values and SNPs with >10% missing values were dropped from the data set resulting in 5316, 5628, and 6231 polymorphic SNPs for the Ki3/NC356, Tx740/NC356, and LH82/LAMA populations, respectively. Crosspoints were predicted using the crosspoint subcommand of SNPbinner [64] to clean data set of double recombinants. The emission probability was set to 0.9 (-p 0.9), the continuous genotype region was set to 0.1% (-r 0.001) of the chromosome size, and the transition probability was calculated using a crosscount of 7,500,000 (-c 7,500,000). The visualize subcommand was used to evaluate the efficiency of the calculated break points to the original SNP calls and identify satisfactory crosspoint parameters. The crosspoint output identified break point locations for each RIL and the prediction of genotypic homogeneity of each region between breakpoint and the SNP calls were adjusted accordingly. Marker datasets filtered by SNPbinner were constructed into linkage maps using the MAP function of QTL IciMapping version 4.1.0.0 (http://www.isbreeding.net/) software. Redundant markers were identified using the “BIN” functionality and redundant markers with greater missing data rate were excluded. Linkage groups were defined by “By Anchor Only” setting and the marker orders were defined by their physical locations using the “By Input” ordering algorithm. Recombination frequencies between markers were calculated based on F_3_ marker frequencies by denoting the “POP.ID” to eight.

The final genetic maps consisted of 1530, 2571, and 2324 SNPs after removal of redundant markers. The genetic map distances were calculated in QTL IcIMappering using the Kosambi mapping function, and the total map lengths were estimated to be 1315, 1207, and 1474 cM for the Tx740xNC356, Ki3xNC356, and LH82xLAMA populations, respectively.

### Linkage Mapping

The entries phenotyped in 2018 were advanced several generations following initial DNA extraction at F_3:4_ and were evaluated in the field at F_6_ generation or greater. For this reason, heterozygous calls (1) were set to missing (−1) and QTL analysis was performed assuming RIL genotype frequencies (“POP.ID” = 4). Analysis by other methods (e.g. treating as F_3_) were also tested to ensure conclusions were similar, but detection power was much lower, likely due to the software trying to fit dominance effects expected to be rare or absent by the F_6_ generation. Inclusive Composite Interval Mapping [65] of Additive (ICIM-ADD) QTL was conducted in the QTL IciMapping v4.1 using the BIP (QTL mapping for bi-parental populations) function. The step parameter was set to 1.0 cM and the probability of inclusion in the stepwise regression (PIN) was set to 0.001. The focus of this study was on understanding the temporal shifts in the marker trait associations of plant height, rather than identifying regions of high confidence that could be used in later marker assisted selection. For these reasons, we defined QTL of interest liberally as those with LOD > 2.0 and percent variation explained ≥ 3% [66], however LOD and other metrics are provided to extract more conservative thresholds. Using the imputed heights from 1 to 85 DAS, ICIM-ADD was performed on each DAS, for each population in each environment separately to access the temporal shifts in allelic effects and marker–trait associations.

A list of candidate genes was obtained from Wallace, Zhang (25). In short, candidate genes were identified from (i) literature, (ii) mining the MaizeGDB database for known height mutants, and (iii) searching the maize genome annotation on Phytozome genes annotated with “auxin”, “brassinosteroid” and/or “gibberellin”. Distance for the center of the QTL confidence interval to nearest candidate gene with the same chromosome were identified.

### Data Availibility

All of the raw and processed data relevant to this study is publically available on Dryad Digital Repository (68). All raw and processed image output files from this study are publicly available and can be obtained by request to the authors.

## Acknowledgements

The authors would like to acknowledge Misty Miles and all members of the Texas A&M UAS project. David Rooney, Jacob Pekar and Stephen Labar for their agronomic and technical support. Graduate students and undergraduate/high school employees for their hard work and effort maintaining fields and collecting phenotypic data. Special thank you to AgReliant Genetics, LLC. and Dr. Ivan D. Barrero Farfan for funding and conducting the genotyping for this research. This project was made possible by financial support from USDA-NIFA-AFRI Award No. 2017-67013-26185 USDA-NIFA Hatch funds Texas A&M AgriLife Research the Texas Corn Producers Board the Iowa Corn Promotion Boardthe Eugene Butler Endowed Chair in Biotechnology and the Texas A&M College of Agriculture and Life Sciences Tom Slick Senior Graduate Fellowship.

## Author contributions

S.L.A. conceptualization, data curation, formal analysis, investigation, methodology, original draft, review and editing (lead), supervision, validation, visualization; S.C.M. conceptualization, funding acquisition, methodology, project administration, supervision, resources, review and editing (supporting); Y.C. data curation, methodology, review and editing (supporting);L.M. data curation, formal analysis, investigation, methodology, review and editing (supporting), software; S.P. conceptualization, funding acquisition, resources, review and editing (supporting), software, supervision; D.C.: conceptualization, funding acquisition, resources, review and editing (supporting), supervision; A.C. data curation, formal analysis, investigation, methodology, review and editing (supporting), software; J.J. data curation, formal analysis, investigation, methodology, resources, review and editing (supporting), software, supervision; I.D.B.F.: data curation, resources, funding acquisition, review and editing (supporting).

## Supporting information

**S1 Fig. Tx740xNC356 correlation heatmaps.** Heat map comparing correlations between manual terminal plant height (PHT), flowering time (DTA/DTS), functional parameters (asymptote, growth rate, inflection point), and UAS P95 estimates by flight date for the Tx740xNC356 population under **[a]** irrigated and **[b]** non-irrigated watering regimens. Growth rate is an empirical constant of the Weibull function which defines the maximum absolute growth rate (m d^-1^).

**S2 Fig. Ki3xNC356 correlation heatmaps.** Heat map comparing correlations between manual terminal plant height (PHT), flowering time (DTA/DTS), functional parameters (asymptote, growth rate, inflection point), and UAS P95 estimates by flight date for the Ki3xNC356 population under [a] irrigated and [b] non-irrigated watering regimens. Growth rate is an empirical constant of the Weibull function which defines the maximum absolute growth rate (m d-1).

**S3 Fig. LH82xLAMA correlation heatmaps.** Heat map comparing correlations between manual terminal plant height (PHT), flowering time (DTA/DTS), functional parameters (asymptote, growth rate, inflection point), and UAS P95 estimates by flight date for the LH82xLAMA population under [a] irrigated and [b] non-irrigated watering regimens. Growth rate is an empirical constant of the Weibull function which defines the maximum absolute growth rate (m d-1).

**S4 Fig. Visual representation of temporal QTL.** Significant temporal height QTL. Red indicates positive allelic effect estimates, blue indicates negative allelic effect estimates, and black indicate non-significant (NS) genomic regions. Lines represent each of 5316, 5628, and 6231 polymorphic SNPs for the Ki3/NC356, Tx740/NC356, and LH82/LAMA populations, respectively, across the genome (X-axis).

**S5 Fig. Visual representation of temporal allele effect estimate**. Visual representation of temporal single marker analysis of the Weibull imputed height estimates. Lines represent each of 5316, 5628, and 6231 polymorphic SNPs for the Ki3/NC356, Tx740/NC356, and LH82/LAMA populations, respectively, across the genome (X-axis).

**S1 Table. Summary of 2018 UAS flight dates.** Summary of 2018 UAS flight dates of the fields containing the Tx740xNC356, Ki3xNC356, and LH82xLAMA populations, including: days after sowing (DAS), the number of images captured, the number of calibrated images, spatial resolution of the mosaic image and mean errors of the GCP geo-referencing.

**S2 Table. Descriptive statistics of UAS flight dates by population.** Summary statistics of the entries for each population (Tx740xNC356, Ki3xNC356, and LH82xLAMA) across the six identified flight dates with high quality point clouds for the irrigated and non-irrigated trials.

**S3 Table. Manual plant height QTL.** Summary of QTL identified using manual terminal plant height as the associated phenotype. Physical locations (bp) based on B73 RefGen_3, AGPv3.

**S4 Table. Flowering time QTL** Summary of significant QTL for flowering time. Physical locations (bp) based on B73 RefGen_3, AGPv3.

**S5 Table. Function growth parameter QTL.** Summary of significant QTL for functional parameters of the Weibull sigmoid function. Physical locations (bp) based on B73 RefGen_3, AGPv3. Growth rate (GR) is an empirical constant of the Weibull function which defines the maximum absolute growth rate (m d^-1^).

**S6 Table. Temporal height QTL.** Summary of significant temporal QTL for height estimates imputed from Weibull sigmoid curve at discrete time points (i.e. DAS where significant associations were identified.). Physical locations (bp) based on B73 RefGen_3, AGPv3.

**S1 File. Statistical methods to identify flight to remove from temporal dataset.**

